# Seasonal Dynamics of Nonstructural Carbon Compounds in Pine Forest

**DOI:** 10.64898/2026.03.05.709835

**Authors:** Clement Kyei Sarpong, Malik Nkrumah, Benju Baniya, Dohee Kim, Asko Noormets

## Abstract

Non-structural carbon compounds (NSCs) serve to buffer short-term imbalances between carbon supply and demand in trees; however, their seasonal dynamics throughout the entire tree remain inadequately understood. We quantified year-round non-structural carbohydrate storage and fluxes in a temperate pine forest by integrating monthly measurements of soluble sugars, starch, and lipids across five tissues with biometric scaling to ecosystem stocks. Soluble sugars were consistently highest in canopy tissues and maintained a relatively stable concentration, even as sugar fluxes exhibited pronounced seasonal variations and reversals. In contrast, starch showed clear seasonality, increasing during the mid-growing season and decreasing later, whereas lipid pools remained relatively stable and contributed minimally to short-term fluctuations. Ecosystem-scale analyses indicated that sugars predominantly contributed to NSC turnover, accounting for approximately 80% of the total annual flux, while stored pools exhibited slower changes. The net annual NSC flux, approximately 65 g C m^−2^ yr^−1^, was relatively modest in comparison to biomass production, which totaled around 522g C m^−2^ yr ^−1^. These findings indicate that seasonal changes in carbon balance are primarily driven by rapid redistribution of soluble carbon rather than by significant changes in overall NSC storage.

## Introduction

Forest ecosystems are significant carbon sinks, sequestering approximately 90% of global carbon biomass(Klein & Hoch, 2015). Forest ecosystem carbon sequestration depends not only on carbon uptake through photosynthesis but also on the allocation of assimilated carbon among respiration, growth, storage, and transfers to soils(Blessing et al., 2015; Guo et al., 2022). In contrast to the storage of structural biomass and the exchange of carbon via photosynthesis and respiration, the temporal and spatial dynamics of nonstructural carbon compounds remain poorly understood (Furze et al., 2018; Miao et al., 2022).

Nonstructural carbon compounds-primary sugars, starch, and lipids—form dynamic pools that buffer short-term imbalances between carbon input and metabolic demand(Dietze et al., 2014). Assimilated carbon initially enters plant tissue as soluble sugars, which may be used directly for respiration and growth or temporarily stored as starch or other compounds(El Omari, 2022; Herrera-Ramirez et al., 2021). These pools support maintenance respiration, tissue turnover, and growth processes, while allowing plants to persist through the periods of reduced photosynthesis. In addition, NSCs contribute to osmotic balance and cellular maintenance during environmental stress(Hui-Ying et al., 2024; Piper, 2021; Zepeda et al., 2022).

According to the conventional theory of nonstructural carbohydrate dynamics in temperate woody plants, seasonal patterns of NSC accumulation and remobilization reflect the balance between carbon supply from photosynthesis and carbon demand for growth, maintenance, and respiration(Furze et al., 2019). During the growing season, when photosynthetic supply exceeds sink demand, particularly as growth slows, NSC reserves accumulate(Prescott et al., 2020), whereas during the dormant season, reduced or absent photosynthesis leads to the mobilization of stored NSCs to sustain respiration and basic metabolic function(El Omari, 2022; Oswald & Aubrey, 2024).

Despite extensive study, whole-tree NSCs storage remains poorly characterized, with limited understanding of how total NSC pool size, seasonal fluctuations, and organ-specific contributions vary among temperate forest trees. Although previous research has estimated overall non-structural carbohydrate storage at the whole-tree level (Hoch et al., 2003; Richardson et al., 2015), comprehensive assessments of total NSC storage throughout the year with high temporal resolution have not been conducted. Obtaining accurate estimates of total NSC storage for the entire tree requires a meticulous approach that involves frequent measurements of NSC concentrations across organs, scaling these measurements to the organ level, and subsequently aggregating them. Such estimates are crucial for understanding the dynamics of carbon flow within trees over time. In this study, we estimate the total NSC in the entire tree of shortleaf pine. Our goal is to track whole-tree NSC dynamics throughout the year and evaluate the proportionality of carbon allocation to biomass growth and non-structural storage, to test the implicit assumption of fixed proportionality in current carbon cycle models.

## Materials and Methods

### Study Site

This study was conducted at the Davy Crockett National Forest (US-CRK), Texas, USA (31°27.774′ N, 95°20.49′ W), which experiences a humid subtropical climate. The forest is dominated by naturally regenerated shortleaf pine (*Pinus echinata*) stands, with few loblolly pine and oaks, managed with biannual prescribed fire. Mean annual air temperature is approximately 19 °C, and mean annual precipitation is 1,148 mm.

### Biomass assessment

Tree DBH was measured monthly in permanent circular plots (radius = 18 m, area = 1,018.3 m^2^) established in January 2021. Four plots were monitored in 2024, with band dendrometers installed on 10 representative trees per plot. Additionally, annual vegetation surveys are done to track annual changes in biomass. Trees were selected based on diameter distribution and species composition in each plot. Band dendrometers (Phytogram, USA; and UMS GmbH, Germany) installed at 1.37 m (DBH) record continuous stem-diameter measurements. Growth was defined as irreversible stem expansion following the zero-growth framework (Zweifel et al., 2016), with growth counted only when *DBH*_*t*_ > *DBH*_*t*−1_. Tree biomass was estimated from DBH using loblolly pine allometric equations (Gonzalez-Benecke et al., 2014) as currently available equations for shortleaf pine do not provide tissue-level biomass estimates(Jenkins et al., 2003). Total aboveground stump biomass (TASB) was calculated as

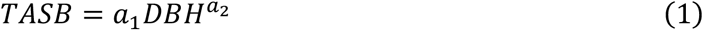

here a1 and a2 are the coefficients. Branch and foliage biomass were calculated as

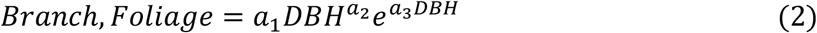

where a1, a2, and a3 are the tissue specific coefficients. Stem biomass was calculated as the difference between TASB and branches and foliage (*Stem* = *TASB* − (*Branch* +*Foliage*)). Coarse-root biomass was estimated according to Jenkins et al. (2003) as:

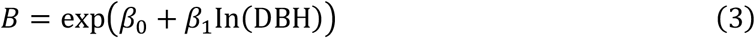

where b0 and b1 are the softwood-specific parameters.

Since dendrometers were installed on a subset of trees within each plot, biomass estimates were scaled to represent the entire plot by employing a basal-area proportion factor.

For each plot:

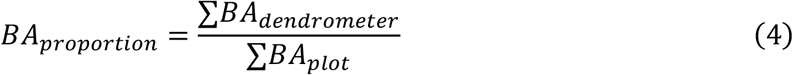

where basal area (m^2^ tree^−1^) was calculated as:

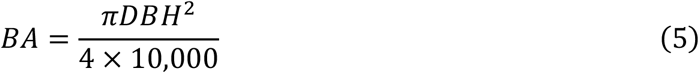

Tree-level biomass totals were divided by this proportion to upscale from sampled trees to entire-plot biomass.

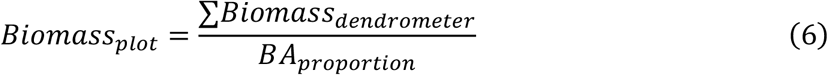

Plot-level biomass (kg plot^−1^) was converted to area-based biomass (kg m^−2^):

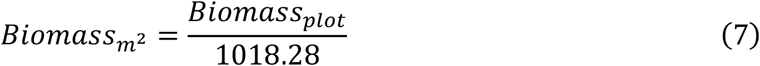

Biomass was converted to carbon, assuming a 50% carbon fraction:

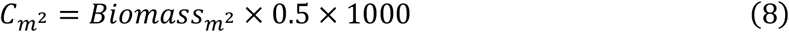

yielding carbon stocks in g C m^−2^.

Monthly biomass production was calculated as the difference in biomass between consecutive months:

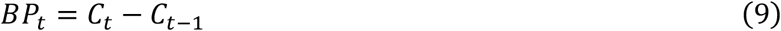

where *C*_*t*_represents biomass (g C m^−2^) at month *t*.

Fine root production was estimated using the bagless ingrowth method. Soil ingrowth volume was prepared by excavating a cylindrical hole (19.5 cm diameter × 30 cm depth), removing all roots and refilling the hole. Cumulative root ingrowth was determined by coring the center of the hole four months later, using a 5 × 30 cm soil corer. For each ingrowth core, dry biomass was converted to carbon, assuming a constant carbon fraction of 0.50. Core-based carbon was then expressed per unit ground area by dividing by the core cross-sectional area:

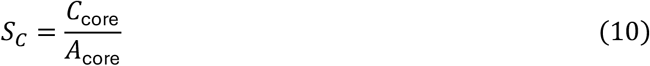

Monthly productivity was calculated by normalizing the incubation length and scaling it to the number of days in the reporting month. For each core:

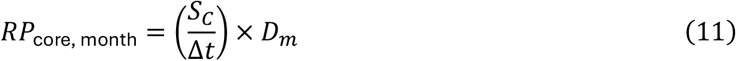

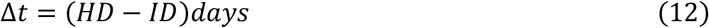

where HD is the harvest date, ID is the installation date, and D_m_ is the number of days in the reporting month

### Plant sampling for NSC analysis

To quantify NSC dynamics, we measured five tissue types (needles, branches, stem wood, coarse roots, and fine roots) and the diameter at breast height (DBH) of 10 trees across four survey plots at a monthly interval throughout 2024.

Leaf and twig samples were collected with a slingshot from sun-exposed mid-canopy. About 10 fascicles of needles and a 10-cm section of branch were collected for each sample. Stem wood was sampled from the north side at about 135 cm height, and coarse roots were sampled with about 50 cm from the base of the tree, both with a 4.3 mm increment borer. Fine roots were sampled with a 5×30-cm soil corer. The samples were stored on ice during the day. Upon returning to the lab at night, the leaves were microwaved at 1000W for 2 min to inactivate the tissue and halt metabolic activity.

Thereafter, all tissue samples were oven-dried at 65 °C for 48 h and finely ground using a ball mill (Retsch MM400). Tissue soluble carbohydrate and starch concentrations were determined according to Sperling et al. (2017), and lipids according to Mishra et al. (2014). Briefly, for each analysis, 25 mg of dry tissue was extracted with 2 mL of deionized water and incubated at 72 °C for 15 min to solubilize free sugars. Samples were then centrifuged at 21,000 g for 10 min, and 200 μL of the supernatant was collected for soluble sugar determination. The remaining pellet was washed with deionized water and gelatinized in 500 μL of 25 mM sodium acetate buffer (pH 4.6) at 100 °C for 1 h. Starch was enzymatically digested by adding 100 μL of amyloglucosidase (70 U mL^−1^; Sigma-Aldrich) and 100 μL of α-amylase (7 U mL^−1^; Sigma-Aldrich), followed by incubation at 50 °C for 2 h to ensure complete hydrolysis. After digestion, samples were centrifuged at 21,000 g for 10 min, and 50 μL of the supernatant was collected for analysis.

### Quantification of Soluble Sugars and Starch

Soluble carbohydrates from both free-sugar extracts and enzymatically digested starch were quantified colorimetrically using the anthrone method, as described by Sperling et al. (2017). Briefly, 50 μL of each sample was diluted tenfold, mixed with 150 μL of anthrone reagent in sulfuric acid (0.1%, w/v), and incubated at 100 °C for 10 min. Absorbance was measured at 620 nm using a microplate reader (SpectraMax, Molecular Devices, San Jose, California). Carbohydrate concentrations were expressed as glucose equivalents.

### Quantification of lipids

Lipid concentration in plant tissues was analyzed using the phosphor-vanillin method as described by(Mishra et al., 2014), with modifications. Ground tissue was saponified in aqueous NaOH for 30 minutes, then acidified with sulfuric acid to liberate free fatty acids. 1 mL of freshly prepared phosphor-vanillin reagent was added to the sample, and the mixture was incubated at 37 °C for 15 minutes to produce a pink color. Absorbance was measured at 530 nm using a spectrophotometer (SpectraMax).

### Ecosystem NSC flux

Ecosystem-level NSC stocks were estimated by scaling tissue-level NSC concentrations with organ-specific biomass production. For each tissue (*j*) and month (*t*), NSC stock was calculated as standing biomass multiplied by the NSC mass fraction, and monthly NSC fluxes were derived from changes in total NSC stock between successive months.

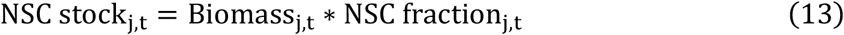

## Results

### Seasonal dynamics of tissue-level NSC concentrations

The tissue-specific concentration data demonstrate a clear and consistent separation of NSC pools throughout the year (Fig. 1). Soluble sugar levels were consistently highest in needles and branches, intermediate in roots, and lowest in stems, showing mild seasonal fluctuations compared to other compounds.

**Fig. 1.**
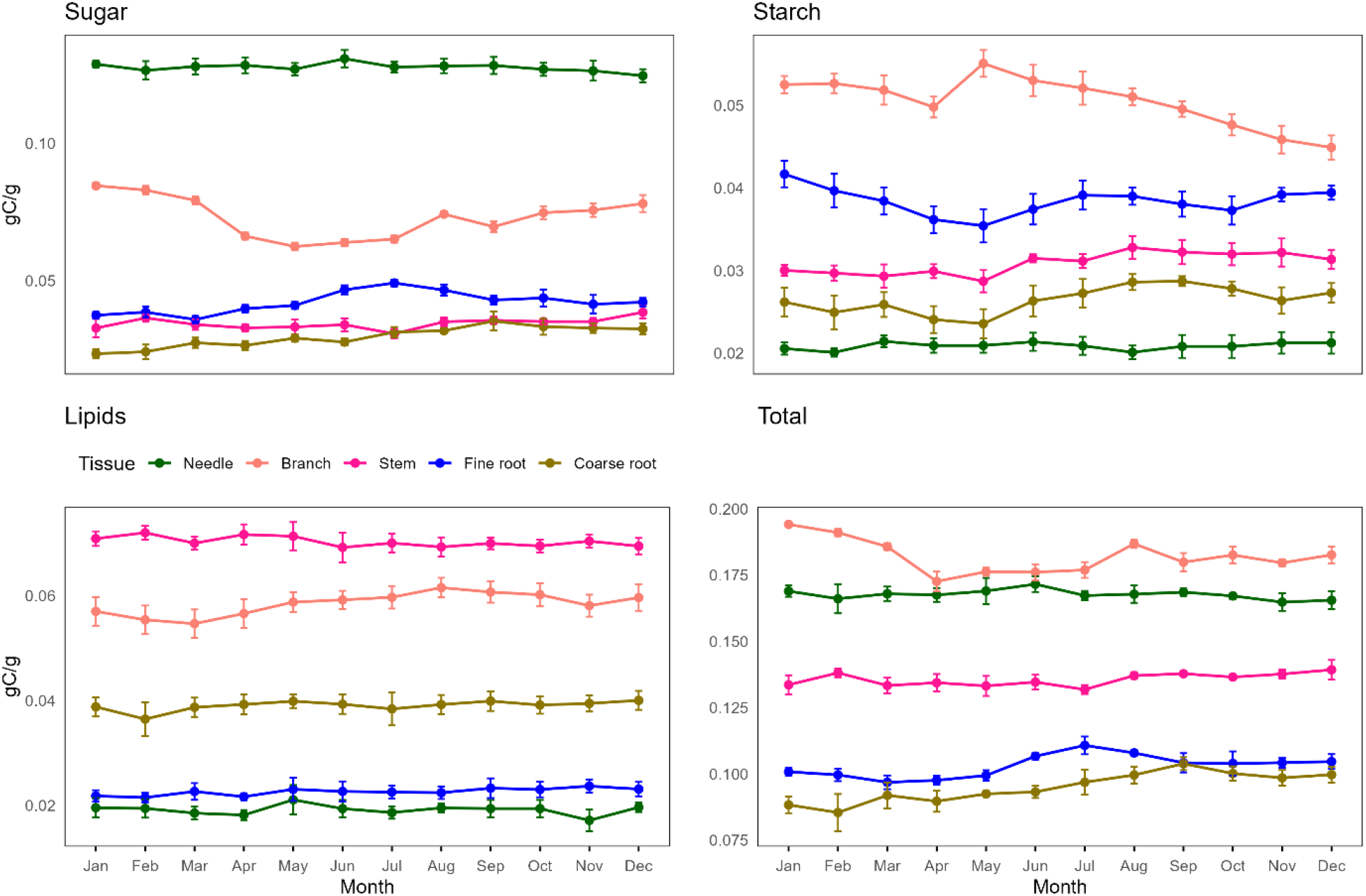
Monthly (mean± SE) concentrations of soluble sugars, starch, lipids, and total nonstructural carbon across five tissues in shortleaf pine in Davy Crockett National Forest in Texas.

**Fig. 2.**
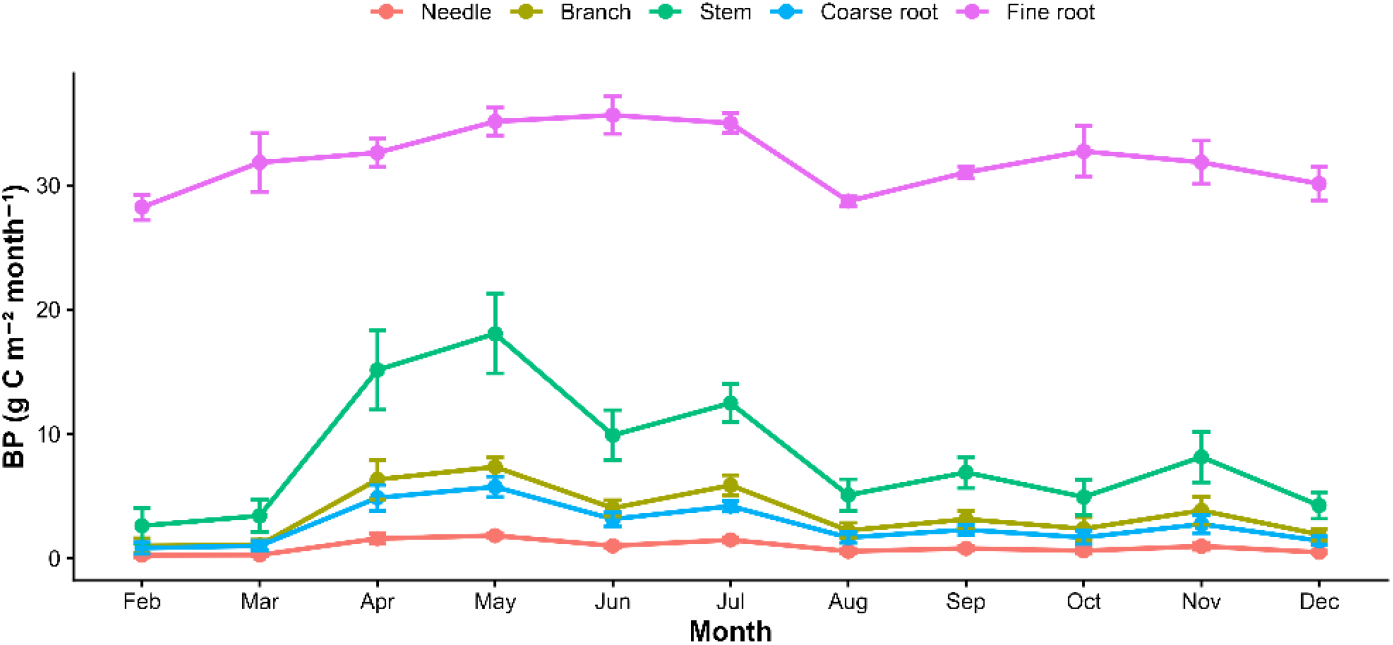
Biomass production (mean± SE) in different tissues of shortleaf pine in Davy Crockett National Forest.

In contrast, starch showed strong seasonality and tissue-specific patterns. Starch accumulated in branches and fine roots during the mid-growth seasons, then gradually declined in late summer and autumn. in needles, the starch concentration is relatively low and stable throughout the year.

Lipid concentrations showed less seasonal variation than starch but remained consistently higher in stems and branches compared to roots and needles.

### Biomass production

Biomass production peaked in April and May in all tissues, except for fine roots, which showed peak growth in May-July. Overall, fine-root productivity was the highest and consistent throughout the year. Other tissues exhibited stronger seasonality but were also all estimated from the same monthly diameter data, whereas fine root biomass and production used a separate coreless ingrowth method. The decline in late-season productivity was associated with water availability.

### Seasonal dynamics of different NSC compounds

The seasonal NSC dynamics were driven primarily by the changes in soluble sugars. Starch showed mid-season accumulation after the growth peaks and small late-season consumption. Lipid concentration was similar in magnitude to that of starch, but its fluxes were largely inversely related to starch.

When grouped into stored NSC pools (starch + lipids) versus sugar (Fig. 3), the temporal dynamics more intuitively mirrored the sink strength of growing tissues (April-May, Aug), secondary growth, and preparation for dormancy. Soluble sugar fluxes showed large and rapid seasonal fluctuations, while stored NSC changed more slowly and with less intensity. Periods of strong sugar mobilization often coincided with near-zero or weakly positive stored NSC fluxes. Conversely, positive stored NSC fluxes during mid-season aligned with decreasing sugar fluxes, indicating that excess carbon was being allocated to storage after meeting immediate metabolic and growth needs.

**Fig. 3.**
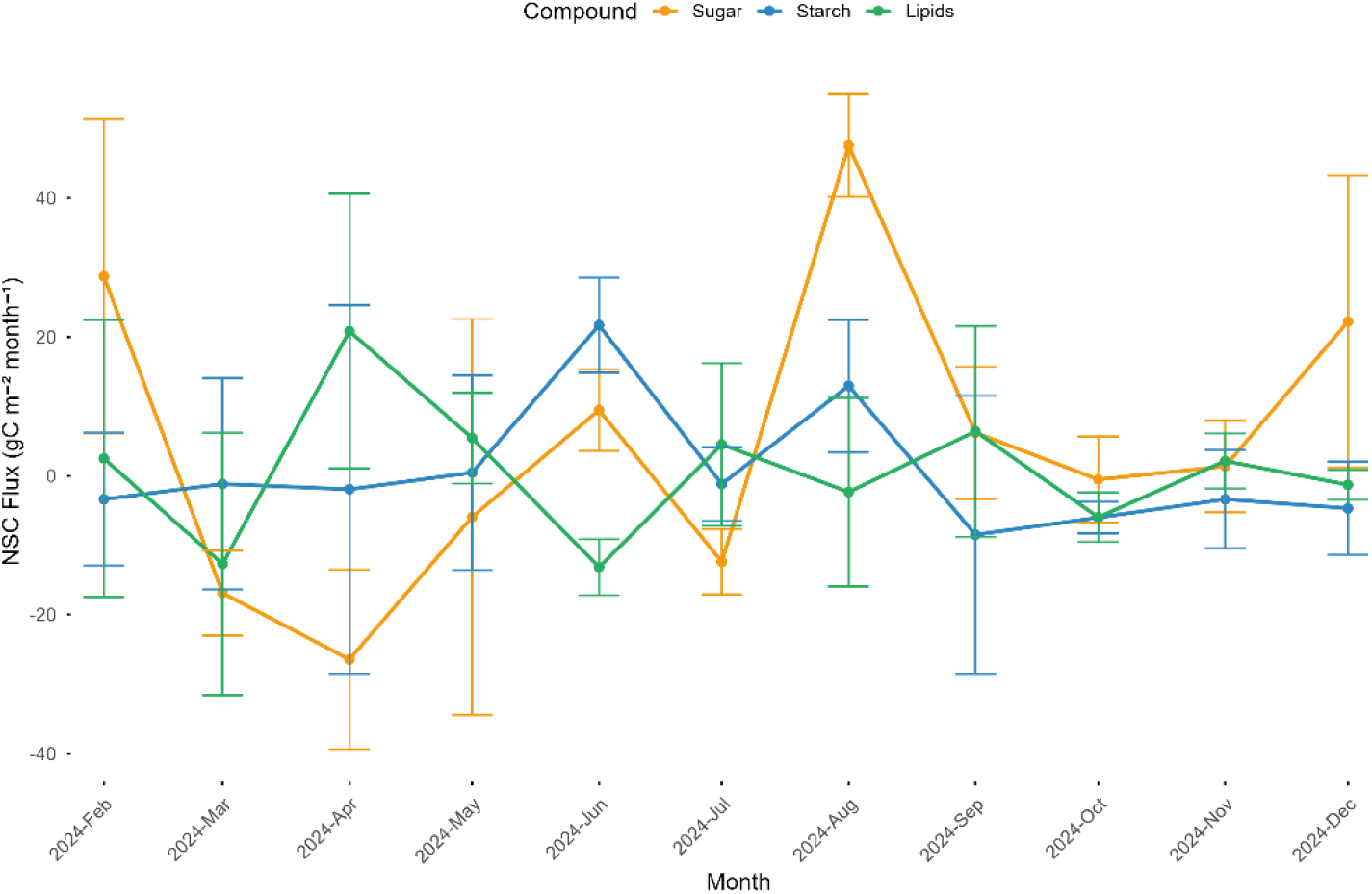
Monthly ecosystem-scale NSC fluxes (mean ± SE) for sugars, starch, and lipids in a shortleaf pine forest in Davy Crockett National Forest in Texas.

**Fig. 4.**
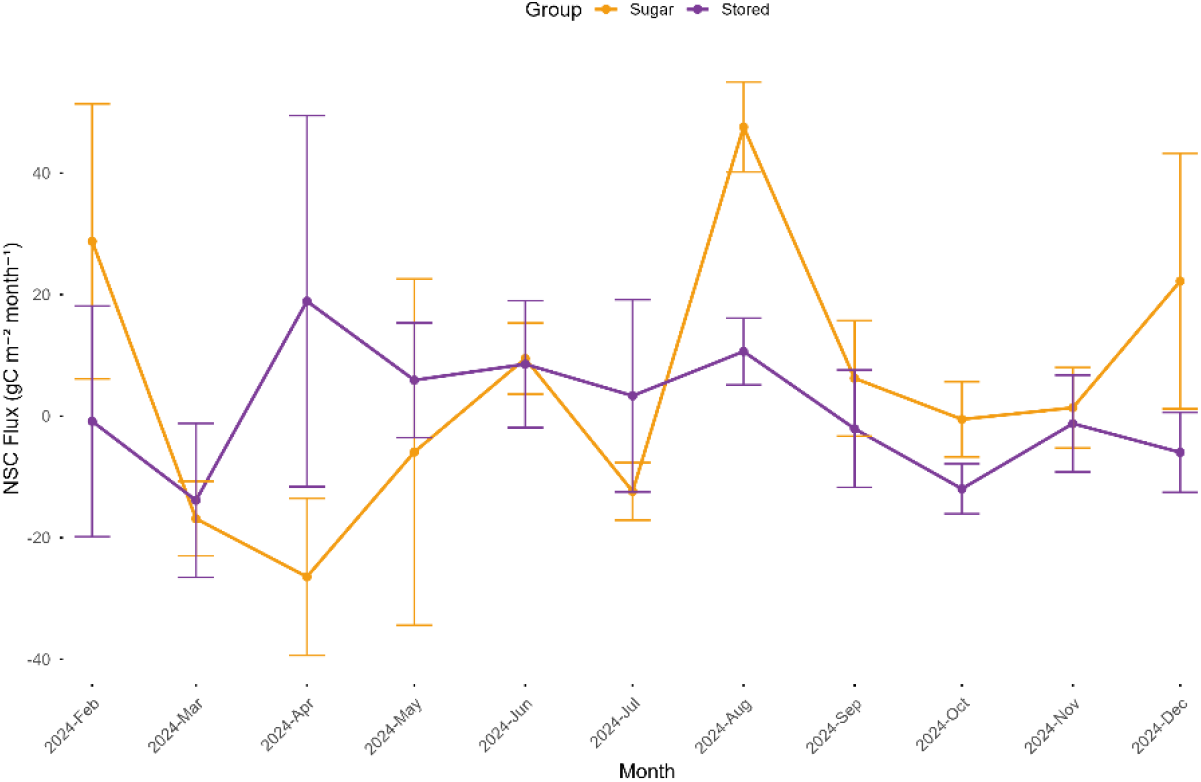
Monthly ecosystem-scale fluxes of soluble and stored NSC (starch + lipids) (mean ± SE).

### Annual NSC budgets and relative contribution to biomass production

The annual integration of ecosystem NSC fluxes captures the overall result of significant seasonal fluctuations in NSC dynamics (Table 1). The total annual NSC flux was 64.92 ±77.03 g C m^−2^ yr^−1^, predominantly influenced by soluble sugars (53.47 ± 47.48 g C m^−2^ yr^−1^). In comparison, starch (4.96 ± 43.46 g C m^−2^ yr^−1^) and lipids (6.49 ± 42.32 g C m^−2^ yr^−1^) contributed minimally to the overall net annual NSC flux. In contrast, the total annual biomass production (522.04 ± 10.63 g C m^−2^ yr^−1^) far surpassed NSC fluxes. Biomass was predominantly allocated belowground, with roughly three-quarters of total production in belowground tissues (382.37 ± 6.18 g C m^−2^ yr^−1^), while aboveground production was 139.67± 9.36 g C m^−2^ yr^−1^.

**Table 1.**
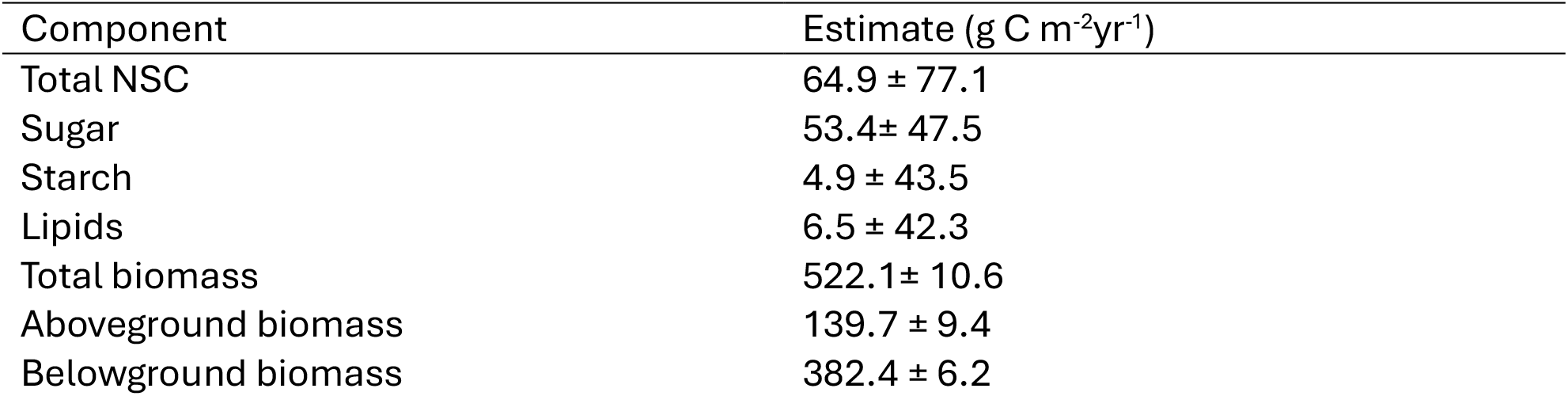
Annual net ecosystem-scale nonstructural carbon and biomass production (mean ± SE). SE represents spatial variability between the 4 survey plots.

| Component | Estimate (g C m<sup>-2</sup>yr<sup>-1</sup>) |
| --- | --- |
| Total NSC | 64.9 ± 77.1 |
| Sugar | 53.4± 47.5 |
| Starch | 4.9 ± 43.5 |
| Lipids | 6.5 ± 42.3 |
| Total biomass | 522.1± 10.6 |
| Aboveground biomass | 139.7 ± 9.4 |
| Belowground biomass | 382.4 ± 6.2 |

## Discussion

This study presents a seasonally detailed, whole-tree evaluation of nonstructural carbon dynamics in a temperate pine forest and demonstrates that seasonal carbon balance is primarily controlled by redistribution among NSC pools rather than by large fluctuations in total NSC storage. By integrating tissue-level concentrations with ecosystem-scale fluxes, our findings refine the traditional view of NSC seasonality and emphasize the importance of flux-based approaches for understanding forest carbon allocation.

During the annual cycle, soluble sugars primarily drove NSC dynamics, making up about 80% of the total NSC flux. These sugar fluxes exhibited significant seasonal variations and often reversed signs, highlighting quick transitions between storage and use. Meanwhile, sugar concentration, especially in needles, remained relatively constant over time, indicating that relying solely on concentration patterns can underestimate true internal carbon dynamics. This supports earlier research indicating that soluble carbohydrates serve as the main short-term carbon currency, buffering mismatches between carbon supply and metabolic demand without needing significant changes in the existing pool size(Dietze et al., 2014; Piper & Tjoelker, 2020).

Stored NSC pools, including starch and lipids, showed a more conservative pattern. Starch accumulated mid-season and was depleted late in the season, reflecting its function as a seasonal reserve rather than a quickly cycled pool. Lipid fluxes were relatively small and more variable, indicating longer residence times and less participation in short-term carbon-dynamics buffering. When combined, stored NSC fluxes were noticeably smoother and smaller in magnitude compared to sugar fluxes. This suggests that short-term seasonal demands are typically satisfied by reallocating soluble carbon rather than significantly depleting storage reserves. Such conservative storage behavior has also been noted in other temperate forest systems and is increasingly seen as evidence of the active regulation of NSC pools(El Omari, 2022; Furze et al., 2019).

These patterns broadly align with traditional models of NSC dynamics, in which carbon accumulates when photosynthetic supply exceeds growth demand and is mobilized when supply declines (Piper & Tjoelker, 2020; Prescott et al., 2020). However, the relatively modest seasonal changes in total NSC storage indicate that surplus carbon is not simply sequestered as long-term reserves. Instead, NSC seems to cycle dynamically, especially through soluble sugar pools, before being partially allocated to starch and lipid reserves.

This interpretation is consistent with the surplus carbon hypothesis, which proposes that growth limitations often constrain carbon use more than carbon assimilation, leading to flexible allocation among transport, storage, and other metabolic pathways (Hoch et al., 2003; Prescott et al., 2020).

From an ecological standpoint, the disconnect between NSC concentrations and fluxes has significant effects on how forests respond to climate variability. As climatic extremes—such as increased vapor pressure deficit and more variable soil moisture—become more common, they are likely to intensify short-term mismatches between carbon supply and demand(Zepeda et al., 2022). In such scenarios, the ability to rapidly mobilize and distribute soluble carbon may be crucial for maintaining respiration and root functions and for recovery after stress, whereas cautious storage strategies may bolster long-term resilience (McDowell et al., 2011; Hartmann et al., 2018)(Hartmann et al., 2020; Hartmann et al., 2015; Hartmann & Trumbore, 2016).

Overall, our findings suggest that seasonal NSC dynamics in pine forests are driven by a coordinated strategy of allocation. This involves the rapid cycling of soluble sugars to meet immediate needs and the conservative storage of pools for long-term security. Such a strategy enables trees to balance short-term metabolic requirements with overall carbon reserves, thereby supporting ecosystem function amid seasonal and climatic fluctuations.

## Conclusion

This study offers a seasonally resolved, comprehensive analysis of nonstructural carbon dynamics within a temperate pine forest, demonstrating that the seasonal carbon balance is primarily influenced by redistribution among NSC pools rather than by significant fluctuations in total carbon storage. Soluble sugars serve as the primary and most flexible form of carbon currency, enabling rapid responses to changing metabolic needs while maintaining relatively stable levels, particularly in canopy tissues. Conversely, starch and lipids serve as stable reserves, showing slower and more restricted seasonal dynamics. The relatively small annual NSC flux compared to biomass production emphasizes the important yet limited role of actively cycling NSC pools. These findings highlight the significance of flux-based methods for understanding carbon distribution and indicate that synchronized soluble carbon cycling, along with conservative storage, sustains seasonal resilience in pine forest ecosystems.

